# First report in France of *Caenoplana decolorata*, a recently described species of alien terrestrial flatworm (Platyhelminthes, Geoplanidae)

**DOI:** 10.1101/2020.11.06.371385

**Authors:** Jean-Lou Justine, Delphine Gey, Jessica Thévenot, Clément Gouraud, Leigh Winsor

## Abstract

Alien land flatworms (family Geoplanidae) are invading many countries in the world. Some can easily be identified by their morphology and colour pattern, but some are more cryptic and necessitate a molecular approach. *Caenoplana decolorata* Mateos et al., 2020 was recently described, from specimens found in Spain, as a sibling species to *C. coerulea* Moseley, 1877. We found that one specimen collected in Nantes, France in 2014 had a 100% identity of its COI sequence with one specimen of the original description of *C. decolorata*, and thus we record here the species for the first time in France.

## Introduction

Alien land flatworms are recorded from many countries worldwide (Sluys, 2016). Some are relatively easy to identify from their morphology and colour pattern. Others belong to species complexes in which the presence of cryptic species make identification by morphology more difficult, necessitating the use of molecular methods. These alien species are generally transported with potted plants, and, as predators of soil animals, can represent a threat to the biodiversity of terrestrial species (Sluys, 2016) and soil ecology (Murchie & Gordon, 2013).

Winsor considered that *Caenoplana coerulea* Moseley, 1877 was a species complex (Winsor, 1997). Alvarez-Presas et al. (2014) studied various specimens from Spain and considered that specimens identified as *“Caenoplana* Ca2” constituted a distinct species (Álvarez-Presas, Mateos, Tudo, Jones, & Riutort, 2014). Recently, these specimens were described as the new species *Caenoplana decolorata* Mateos et al., 2020 (Mateos, Jones, Riutort, & Álvarez-Presas, 2020). The type-material includes a few specimens found in a plant nursery in Girona Province, Spain, in 2012, and no other occurrence has been reported. By its general colour pattern, the species is close to *C. coerulea* Moseley, 1877.

Since 2013, we collected specimens and produced sequence of representatives of the genus *Caenoplana* in France. We checked our sequences to the dataset used by Mateos et al. (2020) and found that one of our sequences matched those of *C. decolorata*, thus showing that the species is present in Metropolitan France.

## Material and Methods

Specimens were collected as a part of a national project in France involving citizen science and molecular characterization of specimens (Justine, Winsor, Gey, Gros, & Thévenot, 2014). We received, from 2013 to now, about 30 specimens of several species of *Caenoplana*. One specimen, now registered in the collection of the Muséum national d’Histoire naturelle in Paris as MNHN JL150, was collected in a glasshouse in the Jardin des Plantes in Nantes, France, during a survey of ants (Gouraud, 2015). The specimen was not photographed when alive, but the collector, one of us (CG), noted that the specimen, found under a pot, was very slender, about 5 cm in length, with a pink head and a yellow dorsal line on a dark background. The specimen was fixed in ethanol where it showed the general morphological feature of *Caenoplana coerulea* in ethanol, i.e. a well-visible dorsal line.

### Molecular analysis

For molecular analysis, a small piece of the body (1-3 mm3) was taken from the lateral edge of the ethanol-fixed individual. Genomic DNA was extracted using the QIAamp DNA Mini Kit (Qiagen). Two sets of primers were used to amplify the COI gene. A fragment of 424 bp was amplified with the primers JB3 (=COI-ASmit1) (forward 5’-TTTTTTGGGCATCCTGAGGTTTAT-3’) and JB4.5 (=COI-ASmit2) (reverse 5’-TAAAGAAAGAACATAATGAAAATG-3’) (Bowles, Blair, & McManus, 1995; Littlewood, Rohde, & Clough, 1997). The PCR reaction was performed in 20 μl, containing 1 ng of DNA, 1× CoralLoad PCR buffer, 3Mm MgCl2, 66 μM of each dNTP, 0.15μM of each primer, and 0.5 units of Taq DNA polymerase (Qiagen). The amplification protocol was: 4’ at 94 °C, followed by 40 cycles of 94 °C for 30’’, 48 °C for 40’’, 72 °C for 50’’, with a final extension at 72°C for 7’. A fragment of 825 bp was amplified with the primers BarS (forward 5’-GTTATGCCTGTAATGATTG-3’) (Álvarez-Presas, Carbayo, Rozas, & Riutort, 2011) and COIR (reverse 5’-CCWGTYARMCCHCCWAYAGTAAA-3’) (Lázaro et al., 2009), following (Mateos, Tudó, Álvarez-Presas, & Riutort, 2013). PCR products were purified and sequenced in both directions on a 3730xl DNA Analyzer 96-capillary sequencer (Applied Biosystems). Results of both analyses were concatenated to obtain a COI sequence of 909 bp in length. Sequences were edited using CodonCode Aligner software (CodonCode Corporation, Dedham, MA, USA), compared to the GenBank database content using BLAST. The final sequence was deposited in GenBank under accession number MW203125.

### Trees and distances

Dr Alvarez-Presas kindly provided the matrix used in the characterization of *C. decolorata* (Mateos et al., 2020). In a preliminary analysis (not shown here), we added all our sequences of *Caenoplana* spp. from France (about 30). We found that some our sequences matched *C. variegata*, some matched *C. coerulea*, and a single one matched the newly described species *C. decolorata*. We then simplified the original matrix: we deleted unnamed species and kept only the sequences which had no indel and were of the same length or longer than our new sequence. The final matrix thus included 14 taxa: 12 sequences from the original dataset (Mateos et al., 2020), an outgroup, *Platydemus manokwari* MT081580 (Gastineau, Lemieux, Turmel, & Justine, 2020), and our single sequence of *C. decolorata*, MW203125.

Using MEGA7 (Kumar, Stecher, & Tamura, 2016), a tree was inferred with the maximum likelihood method, based on the GRT+G model (Nei & Kumar, 2000); all codon positions were used, with 100 bootstrap replications. The study did not intend to provide relationships between species but more simply to check whether our sequences matched those of species characterised in the description of *C. decolorata* (Mateos et al., 2020).

## Results

The tree showed three main branches, each one representing one of the three species of *Caenoplana*, namely *C. variegata, C. coerulea* and *C. decolorata*. The two taxa with several sequences, *C. coerulea* and C*. decolorata*, had 100% bootstrap value, but relationships between taxa showed low bootstrap value and are no more commented here. Our new sequence MW203125 from specimen MNHN JL150 belonged to the *C. decolorata* clade. The similarity between our new sequence and sequence MN990644 was 100% along 781 bp.

## Discussion

Species of *Caenoplana*, which originate from Australia, are often recorded as alien in various countries. *Caenoplana variegata* has been recorded from Italy (Dorigo, Dal Lago, Menchetti, & Sluys, 2020) and Greece (Crete) (Vardinoyannis & Alexandrakis, 2019) (in both cases as *C. bicolor). Caenoplana coerulea* has been recorded from Spain (Álvarez-Presas et al., 2014; Mateos et al., 2013), the Canary Islands (Suárez, Martín, & Naranjo, 2018), the Balearic Islands (Breugelmans, Quintana Cardona, Artois, Jordaens, & Backeljau, 2012), and Argentina (Luis-Negrete, Brusa, & Winsor, 2011).

The identity of species belonging to *Caenoplana* in Europe is now clearer than a few years ago, thanks to several recent works. The presence of *Caenoplana coerulea* has been confirmed with molecular data (Álvarez-Presas et al., 2014). Specimens previously attributed to *C. bicolor* (Graff, 1899) are now attributed to *C. variegata* (Fletcher & Hamilton, 1888) (Jones, Mateos, Riutort, & Alvarez-Presas, 2020). The recent description of the new species *C. decolorata* has added a third binomial species (Mateos et al., 2020).

Our single specimen had a 100% match with one of the sequences of *C. decolorata* from Spain, thus ascertaining that it belongs to the same species. Interestingly, the original description mention that the species was found from beneath pots in a plant nursery, which is exactly where our specimen was found in Nantes, France. Nantes is about 500 km north of the Atlantic Spanish border and about 800 km away from Bordils, the type-locality. However, these distances have no biogeographical signification since the propagation of land flatworms occurs by human transportation of plants and pots. In both cases, specimens were not from the wild but from protected environments, i.e. a plant nursery and a glasshouse. The Spanish specimens were collected in 2012 and our French specimens were collected in 2014; the species is thus present in Europe for at least 8 years, and probably more. For the recently described species *Marionfyfea adventor* Jones & Sluys, 2016, the authors found that the species was already present in several European countries when they became aware of its presence (Jones & Sluys, 2016).

The presence of *C. decolorata* adds one species to the list of alien land flatworms (family Geoplanidae) in Metropolitan France. Currently, the list includes 10 species:

- *Platydemus manokwari*, found only in a hothouse in Caen (Justine et al., 2015; Justine, Winsor, et al., 2014)
- *Bipalium kewense*, widespread in the open, mainly in the South (Justine, Winsor, Gey, Gros, & Thévenot, 2018)
- *Diversibipalium multilineatum*, widespread in the open, mainly in the South (Justine et al., 2018)
- *Diversibipalium* “black”, an undescribed species, found in a single location (Justine et al., 2018)
- *Parakontikia ventrolineata*, with a wide distribution (Justine, Thévenot, & Winsor, 2014)
- *Obama nungara*, with a wide distribution on more than 75% of Metropolitan France (Justine, Winsor, Gey, Gros, & Thévenot, 2020)
- *Marionfyfea adventor*, found in a single location in the wild (Jones & Sluys, 2016)
- *Caenoplana coerulea*, with several locations in the open (Justine, Thévenot, et al., 2014)
- *Caenoplana variegata*, with several locations in the open, locally very abundant (Justine, Thévenot, et al., 2014)
- *Caenoplana decolorata*, found in a single location in a hothouse in Nantes (this paper)

In addition to specimens for which we could obtain sequences, we received a number of records of *C. coerulea* from Metropolitan France based only on photographs obtained by citizen science; since both species are close in colour pattern, it cannot be excluded that some of these are in fact *C. decolorata*.

**Figure 1.**
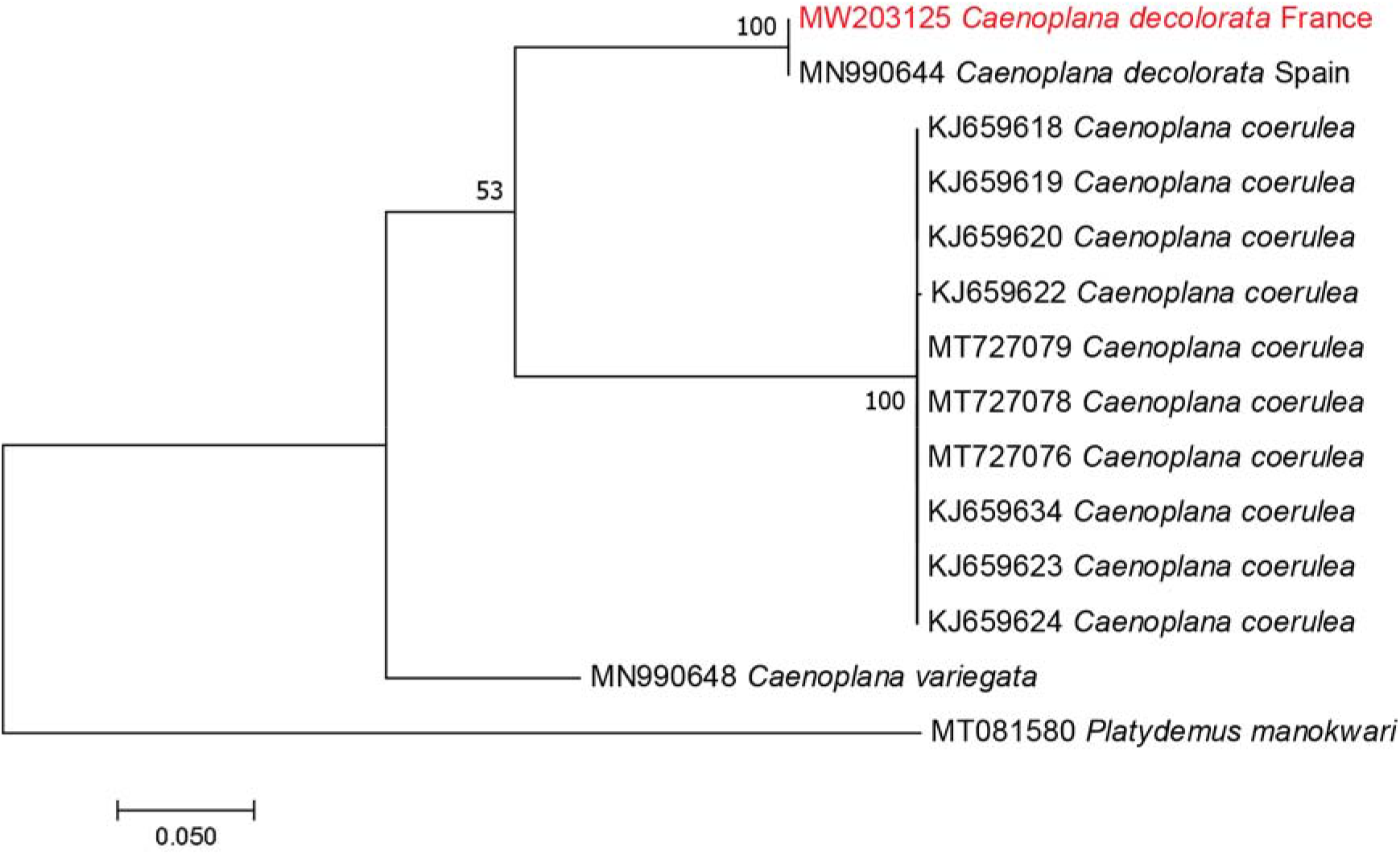
Tree of relationships between species of *Caenoplana* found in France. The matrix was based on that used by Mateos et al. (Mateos et al., 2020), simplified to keep only sequences without indels and of length similar or longer than our new sequence. The evolutionary history was inferred by using the Maximum Likelihood method based on the General Time Reversible model + Gamma distribution. There was a total of 777 positions in the final dataset.

## Acknowledgements

We thank the authors of the description of *C. decolorata*, and especially Marta Álvarez-Presas, for sending us the original matrix used in their paper, thus saving us time and errors retrieving sequences. We thank Philipe Férard, botanist, Jardin des Plantes in Nantes, for allowing the collection of specimens. This work was funded by several Actions Thématiques du Muséum between 2014 and 2020.

